# Metacognition in the audiovisual McGurk illusion: perceptual and causal confidence

**DOI:** 10.1101/2023.03.21.533540

**Authors:** David Meijer, Uta Noppeney

## Abstract

Almost all decisions in everyday life rely on multiple sensory inputs that can come from common or independent causes. These situations invoke perceptual uncertainty about environmental properties and the signals’ causal structure.

Using the audiovisual McGurk illusion this study investigated how observers formed perceptual and causal confidence judgments in multisensory integration tasks under causal uncertainty. Observers were presented with spoken syllables, their corresponding articulatory lip movements, or their congruent and McGurk combinations (e.g. auditory ‘B/P’ with visual ‘G/K’). Observers reported their perceived auditory syllable, the causal structure and confidence for each judgement.

Observers were more accurate and confident on congruent than unisensory trials. Their perceptual and causal confidence were tightly related over trials as predicted by the interactive nature of perceptual and causal inference. Further, observers assigned comparable perceptual and causal confidence to veridical ‘G/K’ percepts on audiovisual congruent trials and their causal and perceptual metamers on McGurk trials (i.e. illusory ‘G/K’ percepts). Thus, observers metacognitively evaluate the integrated audiovisual percept with limited access to the conflicting unisensory stimulus components on McGurk trials.

Collectively, our results suggest that observers form meaningful perceptual and causal confidence judgments about multisensory scenes that are qualitatively consistent with principles of Bayesian causal inference.

## Introduction

Metacognition, the capacity to monitor one’s own uncertainty, is important for adaptive behaviour [1, 2, 3]. In a noisy restaurant, if we feel unsure whether the waiter said ‘beans’ or ‘greens’, we may ask him to repeat it [4-6]. A wealth of research has shown that humans are able to meaningfully report their confidence about perceptual decisions [7-16]. They typically assign a higher confidence to their correct than incorrect decisions. Yet, research to date has almost exclusively focused on simple perceptual decisions based on single cues in one sensory modality, while more naturalistic scenarios such as a busy restaurant expose the brain to numerous signals that may come from common or independent sources. To communicate effectively with the waiter the brain should integrate the waiter’s speech signals selectively with his articulatory lip movements and segregate them from visual and auditory signals produced by other guests. Perception in complex audiovisual scenes thus relies inherently on solving the causal inference or binding problem [17-23].

Bayesian causal inference models (see also Figure 1) address this computational challenge by explicitly modelling the potential causal structures that could have generated the sensory signals. In the case of common sources, signals are integrated weighted by their relative reliabilities into more precise estimates (i.e. fusion). In the case of independent sources, they are processed independently (i.e. segregation) [18, 21]. As the brain does not know a priori whether signals come from common or independent sources, it needs to infer the causal structure from the noisy sensory signals themselves such as them occurring at the same time or location. To account for observers’ uncertainty about the signal’s causal structure, the brain is thought to compute a final estimate by combining the fusion and segregation estimates weighted by the posterior probabilities that signals come from common or independent sources. This decisional strategy is referred to as model averaging (for other decisional strategies see [24]).

**Figure 1.**
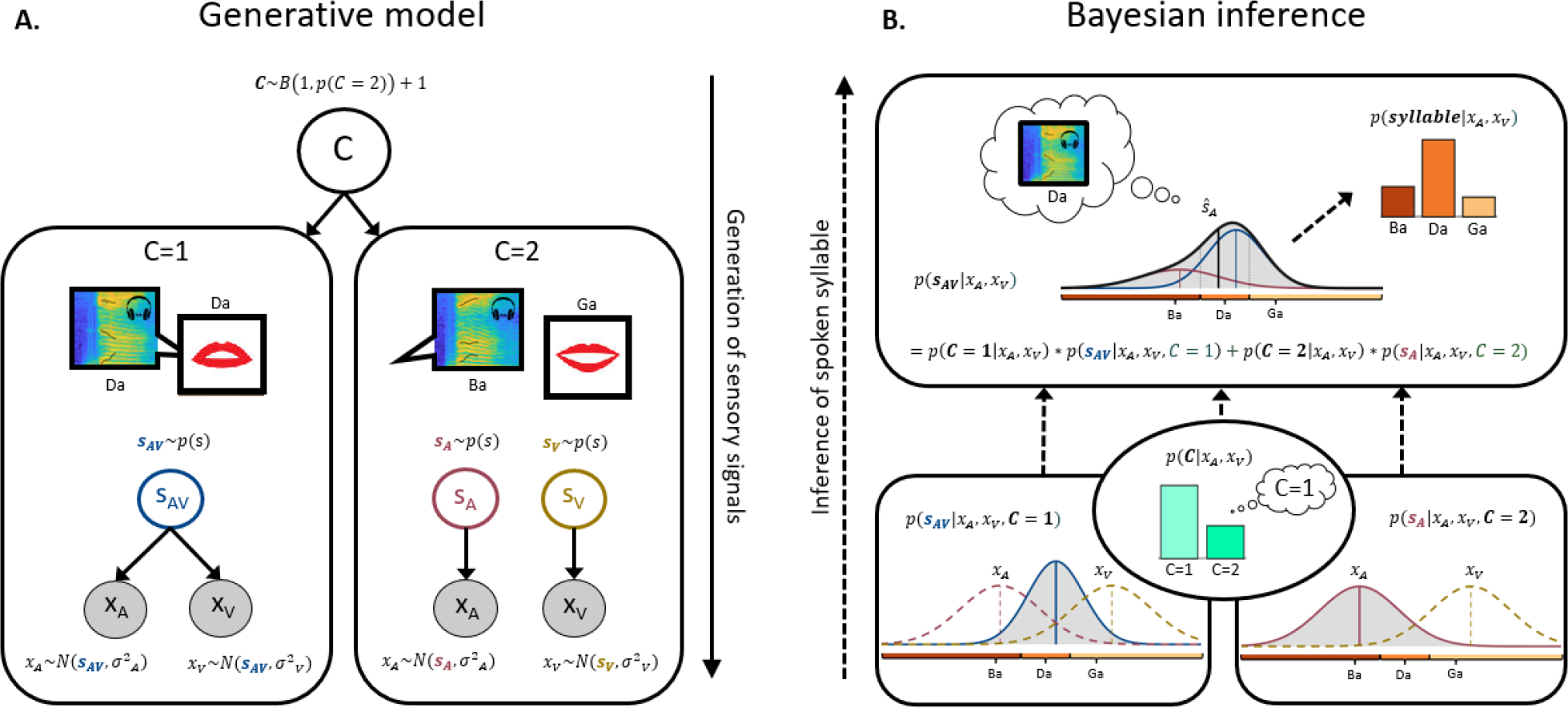
Bayesian Causal Inference model. A. Generative model: The generative model of Bayesian causal inference explicitly models the potential causal structures (C=1 or C=2) that could have generated the auditory and visual signals. A common source (*S*_*AV*_) generates an auditory ‘Da’ speech signal (represented in the figure by a time-frequency spectrogram) and the corresponding articulatory lip movements. Alternatively, an auditory source (*S*_*A*_) generates the ‘Ba’ speech signal and an independent visual source (*S*_*V*_) the lip movements articulating a ‘Da’. The observed auditory and visual signals (*x*_*A*_ and *x*_*V*_) are corrupted by independent noise. B. Bayesian Inference: The observer needs to make two closely related inferences based on the noisy auditory and visual signals: i. perceptual inference about the spoken (i.e. auditory) syllable and ii. causal inference about whether signals come from common or independent sources. In the case of common sources, signals should be fused weighted by their relative reliabilities into a more precise syllable percept (fusion estimate): 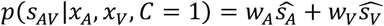 with 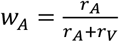 and 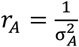. In the case of independent sources, the signal are segregated (segregation estimate). Hence, the phoneme percept should only depend on the speech signals: *p*(*s*_*A*_|*x*_*A*_, *x*_*V*_, *C* = 2). As the observer does not a-priori know whether signals come from common or independent sources, it needs to infer the causal structure from the noisy signals: *p*(*C*|*x*_*a*_, *x*_*V*_). To account for observers’ uncertainty about the signals’ causal structure a final percept is thought to be computed by averaging the fusion and the segregation estimates weighted by the posterior probabilities of common and independent sources. As a result, perceptual and causal estimates arise interactively in the inference process. Adapted from Körding et al. 2007).

Accumulating psychophysics and neuroimaging research has shown that human observers combine sensory signals consistent with the principles of Bayesian causal inference by dynamically encoding segregation, fusion and the final Bayesian causal inference estimates along the cortical hierarchy [25-29]. Yet, little is known about how observers monitor their uncertainties in multisensory environments, in which signals can come from common or separate sources. These more realistic scenarios require the brain to monitor two distinct, but intimately related forms of uncertainty [30]: causal and perceptual uncertainty. Causal uncertainty refers to observers’ uncertainty about the environment’s causal structure, i.e. about whether sensory signals come from common or independent sources. Perceptual uncertainty pertains to observers’ reported perceptual estimate such as their perceived syllable extracted from auditory speech and/or visual facial movements. Causal and perceptual uncertainty interactively arise during perceptual inference and are corrupted by the same sensory noise [18, 21]. At small inter-sensory discrepancies when auditory and visual signals are likely to come from one source, signals are fused into a unified more precise audiovisual percept associated with low causal and perceptual uncertainty. By contrast, at intermediate levels of audiovisual discrepancy observers will be more uncertain about whether signals come from common or independent sources. According to Bayesian causal inference models, the brain would average the fusion and the segregation distributions approximately with equal weight leading to a broader posterior distribution and hence lower perceptual confidence (see Figure 1 and 8 for illustration).

The relationship between causal and perceptual confidence can be studied by explicitly manipulating the discrepancy of the physical stimuli. Yet, even physically identical auditory and visual stimuli may elicit different perceptual and causal decisions because of internal and external noise. This inter-trial variability enables us to characterize the relationship between observers’ causal and perceptual confidence over trials using dual tasks that combine causal and perceptual confidence reports.

This psychophysics study characterized the relationship between perceptual and causal confidence in audiovisual syllable categorization. We presented human observers with spoken syllables (i.e. auditory phonemes), their corresponding articulatory facial movements (i.e. visemes), and their congruent (e.g. visual Ba and auditory Ba) and incongruent (e.g. visual Ga and auditory Ba) combinations. The incongruent phoneme-viseme pairs were designed to elicit ‘B/P’ or illusory ‘D/T’ or ‘G/K’ percepts (i.e. McGurk-MacDonald illusion) [31]. On each audiovisual trial, observers reported their perceived auditory phoneme, the signals’ causal structure and their perceptual and causal confidence.

First, we assessed whether observers integrate audiovisual congruent and McGurk signals into percepts that are associated with greater confidence than their unisensory counterparts. Second, we characterized the complex relationship between causal and perceptual estimates and their associated confidence. Third, we selected audiovisual congruent and McGurk trials on which the sensory signals evoked causal and perceptual metamers, i.e. observers reach the same causal and perceptual decisional outcome. For instance, congruent (i.e. Da-Da) and McGurk (Ba-Ga) trials on which observers report a ‘Da’ percept and a common source are perceptual and causal metamers [30, 32]. We assessed whether despite the same perceptual and causal outcomes observers may still be able to discriminate between them and assign greater levels of causal and perceptual confidence to the congruent than the McGurk trials.

## Methods

### Participants

15 participants were initially recruited, but one did not finish the study. Therefore, 14 participants were included in the analysis (2 males, 2 left handers, mean age: 19.5, range: 18-22). All participants gave written informed consent to participate in this psychophysics study and they were compensated by means of study credits. Participants reported no history of psychiatric or neurological disorders, and no current use of any psychoactive medications. All had normal or corrected to normal vision and reported normal hearing. All participants gave informed written consent to participate in the experiment. The study was approved by the research ethics committee of the University of Birmingham (ERN_11-0470AP4).

### Stimuli

Stimulus material was taken from close-up audiovisual recordings of a female actress’ face on a dark background looking straight into the camera (see Figure 2 A) and uttering the following 18 syllables: ba, be, bi, da, de, di, ga, ge, gi, pa, pe, pi, ta, te, ti, ka, ke, ki. In short, the recordings factorially combined 6 consonants (B, G, D, P, K, T) with 3 vowels (a, e, i). The 6 consonants can be organized into a two-dimensional space spanned by the dimensions of (i) place of articulation (i.e. production place along the vocal tract): B/P (labial), D/T (dental) and G/K (guttural) and (ii) voicing: unvoiced (P, T, K) and voiced (B, D, G). Audio and video were recorded with a camcorder (HVX 200 P; Panasonic). The video was acquired at 25 frames per second phase alternation line (PAL=768×567 pixels); audio was acquired at 44.1 kHz (two channels). The recorded videos were edited (using PiTiVi 0.15.2) into 2000 ms long segments (50 frames) with the first articulatory movement starting at t = 1 second after the beginning of the movie (for further details see [33]). Each video started and finished with a neutral closed lip view of the speaker’s face.

**Figure 2.**
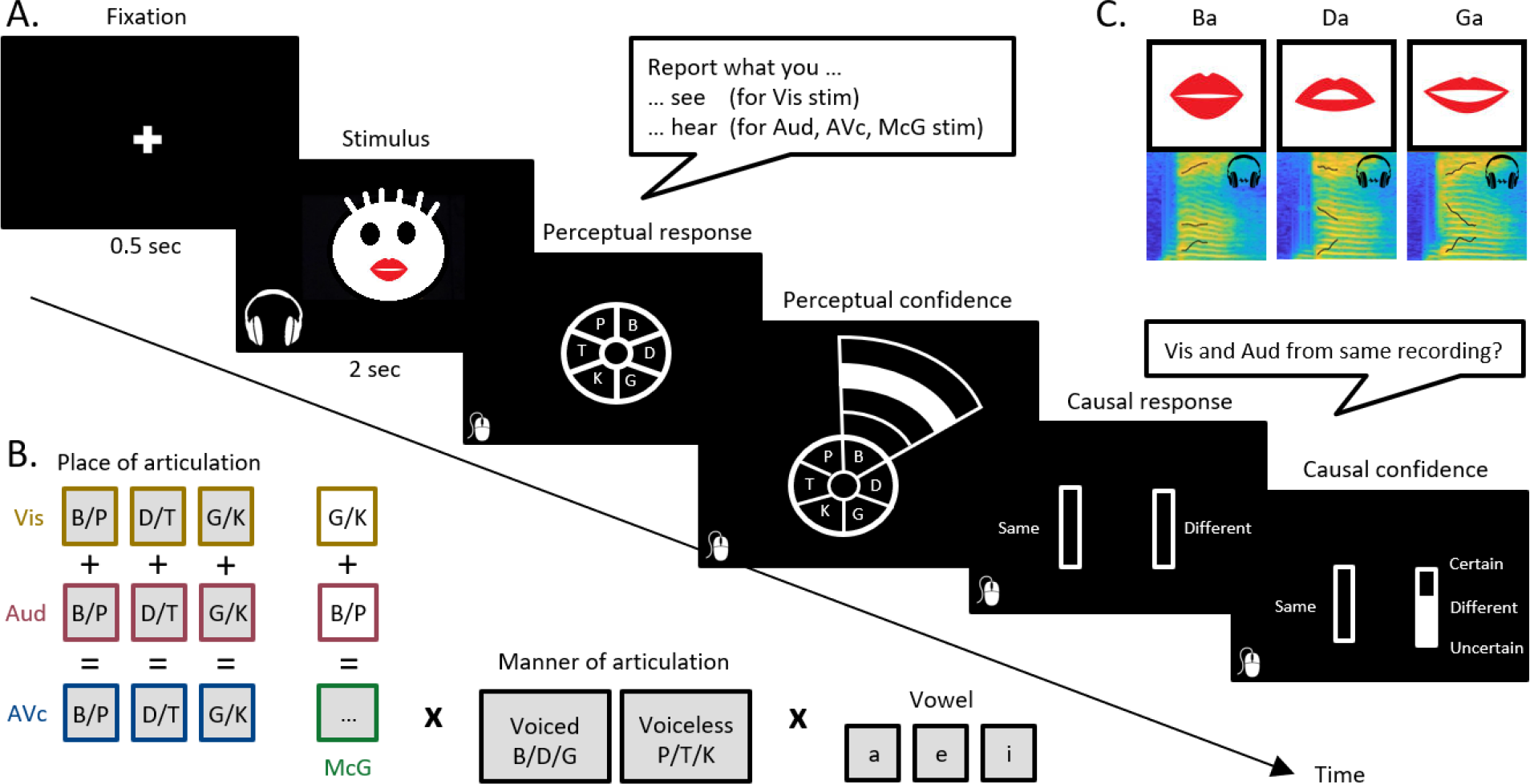
Experimental paradigm, example trial and stimuli. A. Example trial: On each trial observers are presented with a 2 second audio of a spoken syllable (e.g. ‘Ba’), a female face articulating a syllable (e.g. ‘Ba’) or their congruent or conflicting McGurk combinations. N.B. The female face has been changed into an unidentifiable cartoon drawing for this preprint version of the manuscript only. After stimulus presentation, the six possible first letter consonants were presented in a circle. Observers reported the syllable they heard on A and AV trials and the syllable they saw on the V trials. They indicated their perceived first letter consonant by moving their mouse over their preferred response option, after which a layered arc automatically appeared from which participants could select their perceptual confidence level on a 4-point scale (inner layer = low confidence, outer layer = high confidence). On AV trials, observers were then prompted to indicate whether the two auditory and visual stimulus components came from the “same” or from two “different” recordings together with their causal confidence by moving their mouse to a left or right bar. B. Experimental paradigm and stimulus space: Stimuli were videos, audios or their congruent and McGurk combinations. The stimuli differed along place of articulation (labial: B/P, dental: D/T and guttural: G/K), manner of articulation (voiced: B/D/G and unvoiced: P/T/K) and vowel (a, e, i). C. Example stimuli: Video frames of the articulatory positions of a Ba, Da and Ga stimuli and corresponding time-frequency spectrograms of the sounds with the first three formants in black.

We used the movies of 18 different syllables as congruent stimuli. We generated 6 McGurk stimuli by cross-dubbing the video and the audio-track from the B-vowel (auditory) + G-vowel (visual) and the P-vowel (auditory) + K-vowel (visual) stimuli for the three vowels (a, e, i). Further, we presented the corresponding 18 auditory and visual components as unisensory stimuli (see also Figure 2 B).

Finally, we added a considerable amount of white noise to all auditory recordings over the full two-second epoch length. The noise signal was sampled at random from a zero-centred normal distribution with a standard deviation that was equal to 1/3 of the maximum amplitude of the speech signal.

### Experimental Design and Trial

The experiment included AV congruent, AV McGurk trials, unisensory auditory and visual trials. On audiovisual trials, observers reported the first letter consonant that they heard and their perceived causal structure (common vs. independent sources) together with their perceptual and causal confidence respectively. On unisensory trials, observers reported their perceived first letter consonant together with their perceptual confidence.

Each trial started with the presentation of a central fixation cross for 500 ms on a black background. Subsequently, the 2-second stimulus (A / V / AV) was presented. The woman’s upper lip’s screen position approximately matched the location of the fixation cross. On A-only trials, the fixation cross remained on screen during stimulus playback. After stimulus presentation, the six possible first letter consonants were presented in a circle (see Figure 2 A). Observers were instructed to report the syllable they heard on A and AV trials and the syllable that they saw on the V trials. Participants indicated their perceived first letter consonant by moving their mouse over their preferred response option, after which a layered arc automatically appeared from which participants could select their confidence level on a 4-point scale (inner layer = low confidence, outer layer = high confidence). They indicated a response with a left mouse click. Participants could still change their mind by moving their mouse to another letter until they had clicked. This procedure ensured that they provided their perceptual report and associated confidence simultaneously.

On audiovisual trials only, participants were then prompted to indicate whether the two auditory and visual stimulus components came from the “same” or from two “different” recordings by moving their mouse to a left or right bar. The vertical mouse location indicated their causal confidence on a continuous scale (bottom = uncertain, top = certain). Again, participants were allowed to change their minds until they responded by left-mouse button click. Participants were instructed to focus on response accuracy and precision rather than speed. Furthermore, participants were encouraged to scale their confidence responses across trials such that they made use of all 4 levels for perceptual confidence and the entire certainty bar (i.e. from bottom to top) for causal confidence.

The experiment consisted of three 2-hour sessions. It began with a short familiarization block that included 54 trials: 18 AV congruent trials, followed by 18 A-only trials, and finally 18 V-only trials (each syllable appeared once). Before the beginning of each mini-block of 18 trials, the instructions appeared on the screen. For AV and A trials these read “Report what you hear”, whereas for V trials participants were instructed to “Report what you see”.

The main task started after the familiarization block. Participants completed 16 blocks of 144 trials. Each block contained 36 V-only, 36 A-only and 36 AV-congruent trials (2 repetitions of each of the 18 syllable), as well as 36 McGurk stimuli trials (6 repetitions of each of the 6 McGurk stimuli sets). The AV-congruent and McGurk stimuli were randomly interleaved, whereas the unisensory trials were presented in separate mini-blocks.

### Experimental procedure

Participants were seated in a small dark room. They placed their chin on a chinrest that was positioned at a distance of 55 cm from a computer monitor (53 by 30 cm). Visual stimuli were shown at a frame-rate of 25 Hz in a central square of 20 by 20 cm, surrounded by a black background. Auditory stimuli were presented at 44.1 kHz by means of headphones (Sennheiser HD 280 Pro) at a comfortable listening volume (65 dB).

Tobii EyeX eye tracker was used during the experiment to monitor whether participants focused their eye gaze on the centre of screen (+/- 5 degrees) during stimulus presentation. The central area of focus corresponded approximately to the woman’s upper lip. Participants who did not follow task instructions received corrective feedback. The eye tracking data was not further analysed.

The experiment consisted of 2-hour sessions that were performed on separate days, with maximally one week in between successive sessions. Participants had to finish the full experiment within two weeks. Participants were encouraged to take self-paced breaks between blocks within a session.

The experiments were run in Matlab 2014b using the Psychophysics Toolbox 3 [34, 35] and the Tobii Eyex toolkit [36].

### Confidence normalization, statistical analyses, and simulations

#### Confidence normalization

The perceptual and causal confidence distributions varied substantially in mean and spread across participants. We therefore normalized the confidence distributions with the help of the cumulative distribution function. More specifically, each raw confidence value x was mapped onto a normalized confidence value equal to the corresponding value in the cumulative distribution. In cases with several identical raw confidence values, we assigned the average across their normalized confidence values to those:

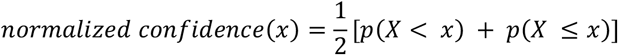

This transformation ensured that the mean over confidence values in all participants was equal to 0.5 and that the confidence values were spread across the entire range. For instance, if a participant indicated maximal causal confidence (top of the bar) in 30% of the trials, the normalized causal confidence values for these trials would all be equal to the mean of 0.7 and 1 = 0.85 (see supplementary Figure S1).

#### Statistical analysis

For each participant we computed the response fractions over the six first letter categories and the four perceptual confidence levels, response accuracies for unisensory and AV congruent conditions and across-trial’s mean perceptual and causal confidence values (after normalization, see above). Because these indices were fractions or probabilities i.e. limited to a range between zero and one, we transformed them using the arcsine-square-root transformation, 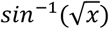, before entering them into general linear mixed effects models (e.g. repeated measures ANOVAs). Greenhouse-Geisser corrections were applied when Mauchly’s test indicated that the sphericity assumption was violated. Statistical analyses were performed with JASP 0.16.3.0 (https://jasp-stats.org/).

#### Simulations

To illustrate the relationship between perceptual and causal decisions as well as their corresponding confidence levels we performed simulations based on the Bayesian causal inference model (for details, see [18]). Originally, the Bayesian causal inference model has been developed to model spatial categorization responses, in which continuous spatial estimates are mapped onto discrete spatial choices. Likewise, our simulations made the assumption that Ba, Da and Ga stimuli lie on a continuous abstract ‘place of articulation’ dimension that goes from labial (i.e. lip) ‘Ba’ to dental (i.e. ‘teeth’) ‘Da’ and finally to guttural (i.e. throat) ‘Ga’. Further, we assumed that this dimension is shared across the visual and auditory senses.

These continuous estimates are mapped onto ‘Ba’, ‘Da’ and ‘Ga’ categories via categorical perception. Auditory and visual senses provide information about the place of articulation via formants for different phonemes and articulatory movements (i.e. viseme). The 2nd formant in the time-frequency spectrograms discriminates between Ba, Da and Ga (see Figure 2 C). Likewise, the articulatory lip movements inform about the place of articulation.

For the example shown in Figure 1 and each of the four examples in Figure 8, we sampled one auditory and one visual signal from 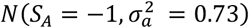 and 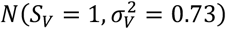 and we computed the likelihoods ℒ(*S*; *x*_*V*_) and ℒ(*S*; *x*_*A*_) (pink and brown dashed lines). Assuming a flat (i.e. uninformative) prior over the place of articulation dimension and a causal prior *P*(*C* = 1) = 0.5, we computed the posterior distribution:

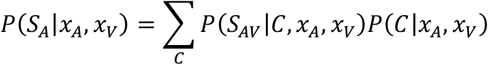

This posterior distribution (solid black line) is a mixture of the full segregation *P*(*S*_*A*_|*x*_*A*_, *x*_*V*_, *C* = 2) (pink solid) and the fusion distributions *P*(*S*_*A*_|*x*_*A*_, *x*_*V*_, *C* = 1) (blue solid) weighted by the posterior probabilities over common and independent causes *P*(*C*|*x*_*A*_, *x*_*V*_) (green bar plots). To obtain the discrete posterior probabilities over syllable categories B/P, D/T and G/K (orange bar plots), we integrated the continuous posterior probability distribution limited by the category boundaries that separate the three response choices (i.e. category boundaries ‘Ba’ vs. ‘Da’ was set to -0.5 and ‘Da’ vs. ‘Ga’ to 0.5). We present these simulation results to provide a qualitative explanation for the pattern of findings in our study. We note that for more complex abstract dimensions such as ‘place of articulation’ several assumptions of the Bayesian model may not fully hold, so that we refrain from formal quantitative modelling (e.g. Gaussian distributions, common decision dimension shared between auditory and visual senses, additional variability within categories etc., for related approaches see [22, 37]).

## Results

In the following we present the key results organized in line with the results figures and supplementary tables. The complete statistical results are presented only in the tables (see supplementary material).

### Performance accuracy and perceptual confidence in auditory, visual and audiovisual congruent conditions (*Figure 3 and Table 1*)

Consistent with Bayesian models of multisensory perception observers integrated audiovisual signals into more precise estimates as indicated by their greater perceptual accuracy. Moreover, because the variance of the posterior distribution of the audiovisual distribution is smaller or equal to that of either unisensory posterior distribution, we would expect participants to be more confident on audiovisual congruent than unisensory conditions. Indeed, in line with these predictions, we observed a significant increase in perceptual accuracy and confidence for the audiovisual congruent conditions (Figure 3 A-B and Table 1 A-B).

**Figure 3.**
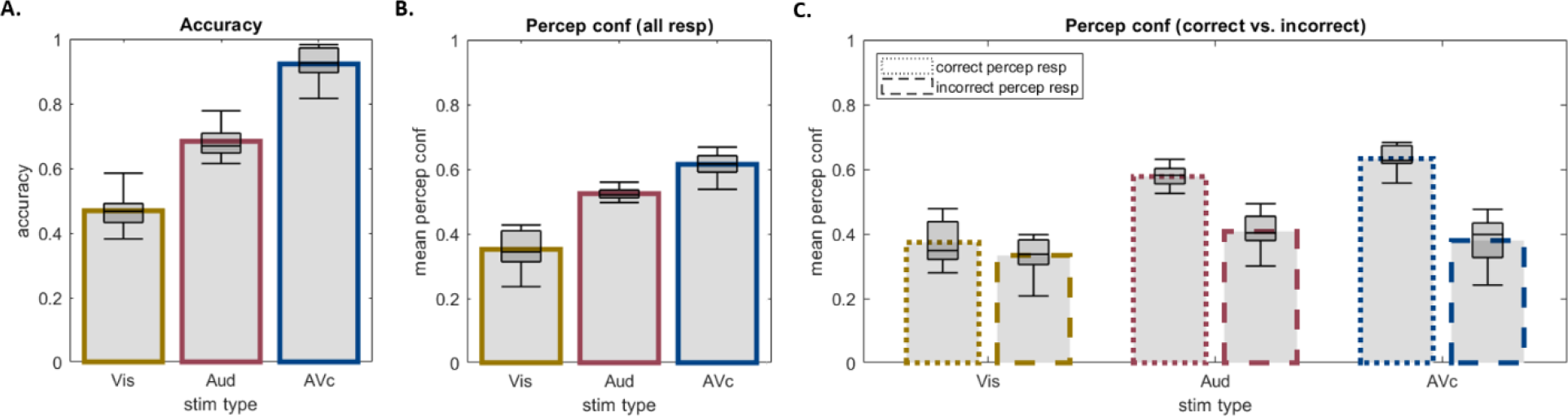
Perceptual accuracy and confidence for auditory, visual, and audiovisual congruent stimuli. A. Perceptual accuracy for V, A and AV congruent: Perceptual accuracy is greater for AV congruent than unisensory A or V stimuli. B. Perceptual confidence for V, A and AV congruent: Perceptual confidence is greater for AV congruent than unisensory A or V stimuli. C. Perceptual confidence for V, A and AV congruent separately for correct and incorrect syllable percepts: We observe an interaction between correct vs. incorrect and sensory modality (A, V AV). In all figure components bar plots show across subjects’ means and box plots indicate across subjects’ median response fraction with first and third quartiles and with whiskers indicating the minimum and maximum subject values.

Moreover, observers were sensitive to their own accuracy as indicated by a significant increase in perceptual confidence for correct relative to incorrect responses (Figure 3 C and Table 1 C). This metacognitive sensitivity was significantly greater for congruent than auditory and in particular than visual conditions (i.e. significant interaction between correct/incorrect x sensory modality). This effect that can be explained by the differences in performance accuracy across sensory modalities. At first sight, it may be surprising that observers were 50% accurate (i.e. better than chance level at 1/6) on the visual first letter categorization task, but showed no metacognitive sensitivity. In other words, they were unable to discriminate between their correct and wrong visual decisions. This metacognitive insensitivity arises because the 6 response options are spanned by the place of articulation (e.g. B vs. D vs. G) and the voicing (i.e. B vs. P) dimensions. While the visual modality is very informative about the place articulation, it is uninformative about the voicing dimension. Hence, the vast majority of errors are ‘voicing errors’ such as misclassifying a ‘Ba’ as a ‘Pa’ stimulus. From the perspective of the voicing dimension, 50% accuracy is equivalent to chance performance. Observers’ incapacity to discriminate between voiced and unvoiced syllables based on visual input alone thus explains their insensitivity to metacognitively discriminate between correct and incorrect visual responses. (See also supplementary Figure S2).

### Perceptual decisions and confidence on unisensory and McGurk trials (*Figure 4 and Table 2*)

These bar plots show how the brain combines auditory ‘B/P’ and visual ‘G/K’ information McGurk conflict trials and their associated confidence levels. The perceptual accuracy for place of articulation is comparable across auditory ‘B/P’ and visual ‘G/K’ trials (i.e. no significant difference, see Table 2 A). Yet, auditory ‘B/P’ stimuli are associated with significantly greater response entropy, i.e. their responses are spreaded across all categories along the place of articulation dimension (Table 2 B). On 10% of the trials an auditory ‘B/P stimulus is perceived as D/T and even G/K. By contrast, a visual G/K stimulus is perceived as a D/T, but nearly never as a B/P syllable. This difference between auditory and visual entropy explains that McGurk stimuli are mainly perceived as D/T and G/K stimuli, i.e. perceptual categories that are consistent with the visual G/K and the auditory B/P stimulus.

**Figure 4.**
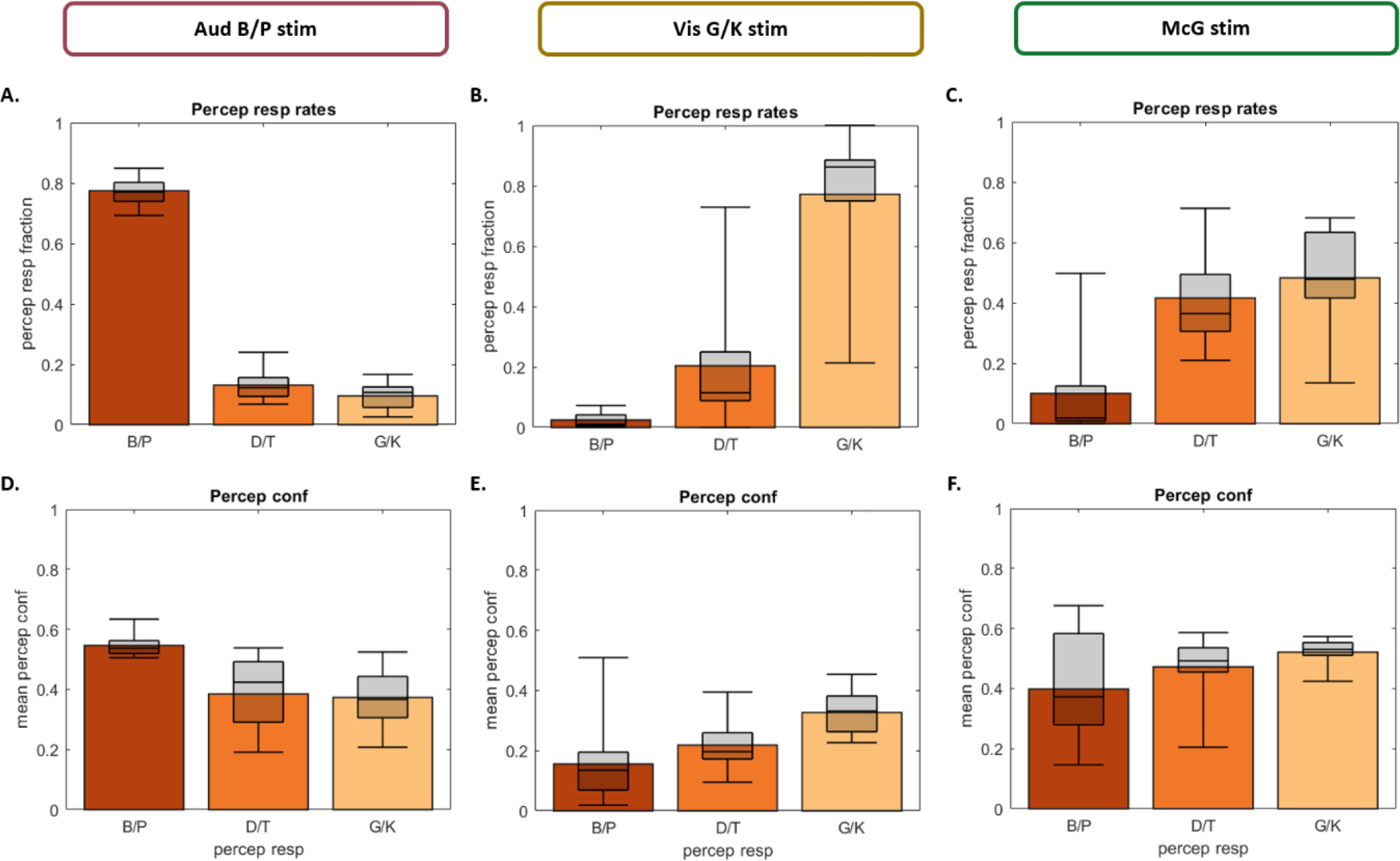
Perceptual response fractions (A, B, C) and perceptual confidence (D, E, F) for auditory B/P, visual G/K and McGurk stimuli. Response fractions over ‘B/P’, ‘D/T’ and G/K’ percepts (top) and associated perceptual confidence (bottom) separately for auditory B/P stimulus (A, D), visual G/K stimulus (B, E) and McGurk stimulus (i.e. auditory B/P and visual G/K; C, F). In all figure components bar plots show across subjects’ means and box plots indicate across subjects’ median response fraction with first and third quartiles and with whiskers indicating the minimum and maximum subject values.

Further, the perceptual confidence is significantly higher for McGurk trials with D/T and G/K outcomes than for unisensory auditory and visual trials (Table 2 D-E), because auditory and visual channels provide partly complementary information about syllable category (see discussion above about voicing and place of articulation in previous section). By contrast, the confidence for a B/P percept is also significantly lower for the McGurk than the unisensory auditory stimuli (Table 2 C). This reflects the fact that on the very few McGurk trials with B/P outcome observers perceive auditory and visual signals as coming from different sources.

### The relationship between perceptual and causal inference (*Figure 5 and Table 3*)

Perceptual and causal decisions are intimately related in the inference process and susceptible to shared sensory noise. These dependencies explain that correct common cause responses on congruent trials go together with correct perceptual responses (Figure 5 A; Table 3 A). Likewise, observers associated a greater causal confidence to correct common cause responses mainly when they categorized the first letter correctly (i.e. significant interaction in causal confidence for common vs. independent cause responses x perceptual accuracy, Table 3 C, Figure 5 C).

**Figure 5.**
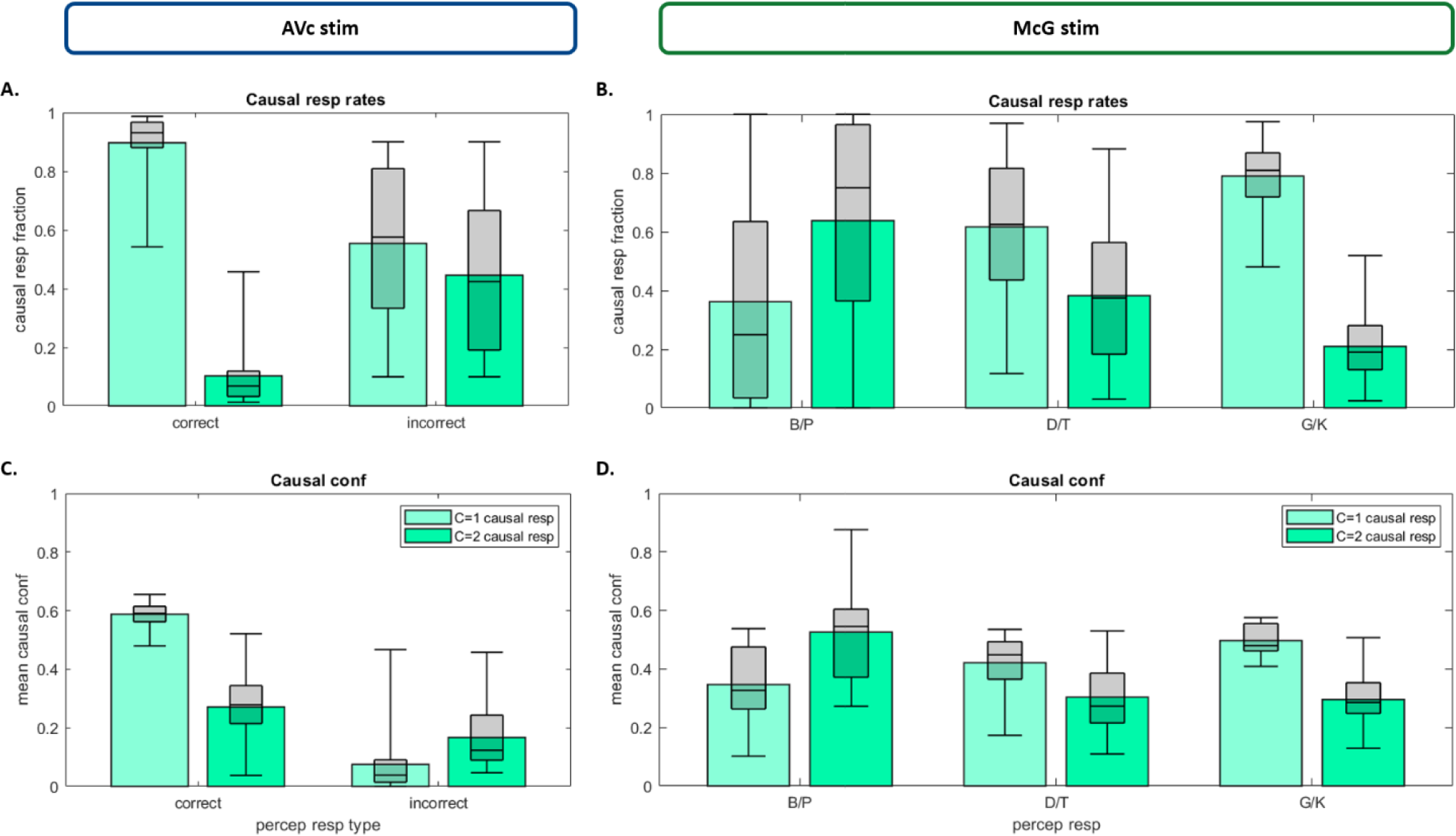
Causal response fractions (A, B) and causal confidence (C, D) for audiovisual congruent and McGurk stimuli. A, C. Response fractions over ‘Common’ (C=1) and ‘Separate’ (C=2) cause responses (A) and the associated causal confidence (C) separately for audiovisual congruent trials on which observers categorized the syllable correctly or incorrectly. B, D. Response fractions over ‘Common’ and ‘Separate’ cause responses (B) and the associated causal confidence (D) separately for McGurk trials with ‘B/P’, ‘D/T’ and ‘G/K’ percepts. In all figure components bar plots show across subjects’ means and box plots indicate across subjects’ median response fraction with first and third quartiles and with whiskers indicating the minimum and maximum subject values.

The close relationship between causal and perceptual inference is also manifest in the McGurk trials. The fraction of common cause responses is directly associated with observers’ perceptual categorization responses, increasing from B/P to D/T and G/K responses (i.e. significant main effect of perceptual category, Table 3 B, Figure 5 B). Likewise, the confidence for common cause responses was greatest for G/K percepts, while the confidence for independent cause responses peaked for B/P percepts (i.e. significant interaction between causal and perceptual outcomes, Table 3 D, Figure 5 D). In other words, as expected, illusory McGurk ‘D/T’ and ‘G/K’ percepts are associated with observers’ decisions and confidence that audiovisual signals share a common cause and should hence be integrated.

### The relationship between perceptual and causal confidence (*Figure 6 and Table 4*)

Our dual task design enables us to characterize the relationship between causal and perceptual confidence based on inter-trial variability. Thus, we sorted observers’ perceptual decisions and confidence according to the causal decision and confidence on those trials (median split per response category for each subject). Consistent with Bayesian causal inference models, common source decisions and high causal confidence were associated with high perceptual accuracy (Figure 6 A) and perceptual confidence (Figure 6 C). Statistically, this observation is supported by significant main effects of causal decision and causal confidence level on observers’ perceptual accuracy and perceptual confidence (Tables 4 A and 4 D).

**Figure 6.**
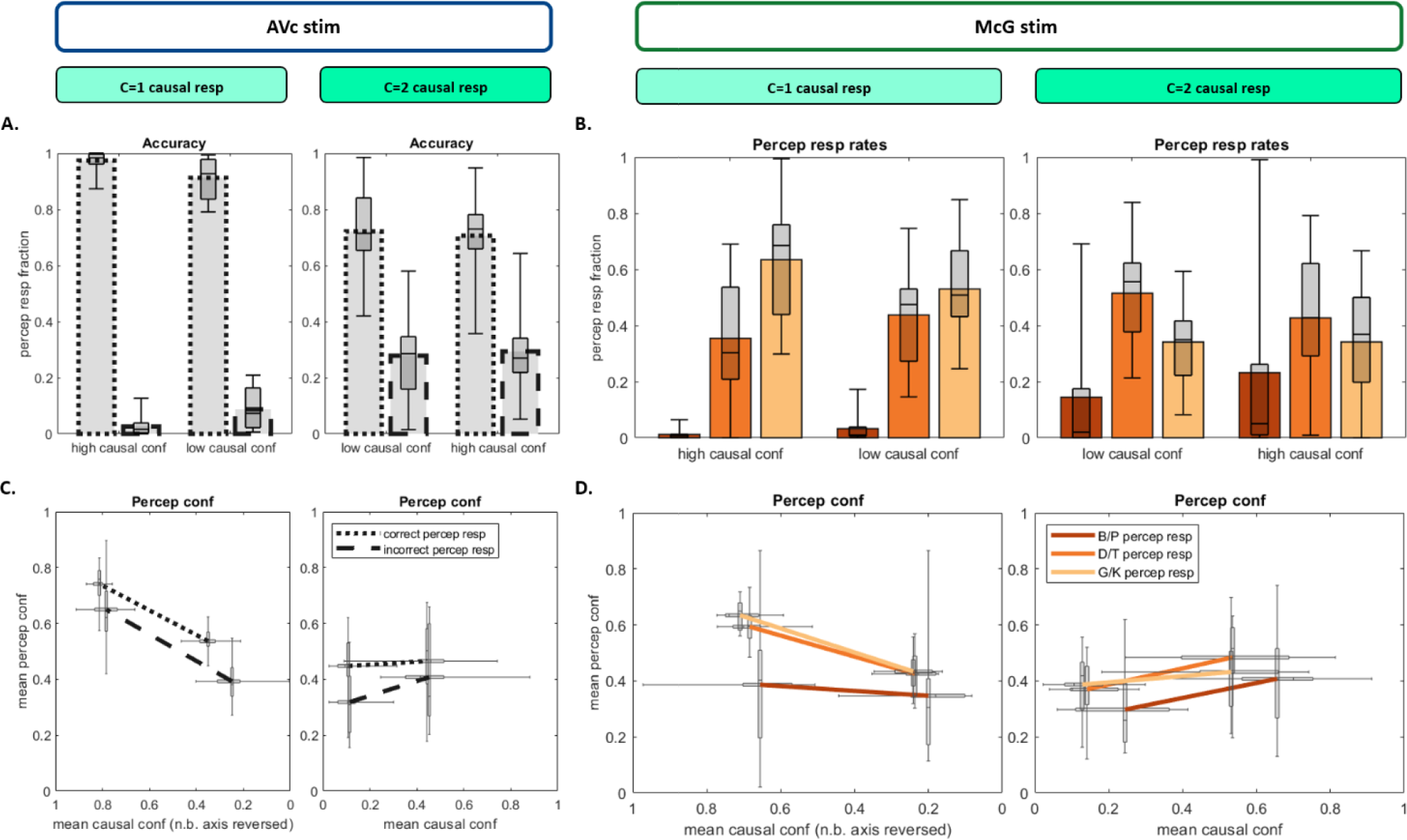
Perceptual and causal response fractions and their associated perceptual and causal confidence. A, C. Fraction of correct and incorrect perceptual responses (A) as well as their associated causal and perceptual confidence (C) separately for ‘Common’ (C=1) and ‘Separate’ (C=2) cause response trials. B, D. Fraction of ‘B/P’, ‘D/T’ and ‘G/K’ percepts (B) as well as their associated causal and perceptual confidence (D) separately for ‘Common’ and ‘Separate’ cause response trials. In all figure components bar plots show across subjects’ means and box plots indicate across subjects’ median response fraction with first and third quartiles and with whiskers indicating the minimum and maximum subject values. Likewise, the crosses in the line plots (C, D) indicate first and third quartiles and the longer cross whiskers the minimum and maximum subject values.

Likewise, on McGurk trials common cause decisions and confidence are closely related with observers’ perceptual outcome and perceptual confidence. As predicted by the Bayesian causal inference model, illusory G/K percepts on McGurk trials were significantly more frequent for common source responses with high causal confidence (i.e. significant main effects of causal decision and post-hoc test for causal confidence on C=1 responses, Table 4 C, Figure 6 B). By contrast, B/P percepts were associated mainly with independent cause decision at high level of confidence (Table 4 B). Furthermore, perceptual confidence was larger when causal confidence was high and it was larger for D/T and G/K percepts than for B/P percepts (significant main effects of causal decision and perceptual response, Table 4 E, Figure 6 D).

In summary, both congruent and McGurk trials demonstrate that causal inference and causal confidence implicitly affect observers’ perceptual decisions and confidence.

### Causal and perceptual metamers (*Figure 7 and Table 5*)

The results presented so far show that on a large percentage of McGurk trials observers’ integrate the auditory ‘B/P’ and a visual ‘G/K’ stimulus into an auditory ‘Da’ or ‘Ga’ percept and report a common cause. On these trials, observers thus integrate audiovisual McGurk signals into perceptual and causal metamers of the corresponding ‘G/K’ and ‘D/T’ congruent trials. We next investigated whether, despite identical perceptual and causal decisions, observers were metacognitively aware that the ‘Da’ and ‘Ga’ percepts on McGurk trials rely on incongruent audiovisual information and hence report lower perceptual and causal confidence values. To address this question, we categorized congruent trials (with correct syllable category and C=1 causal response) and McGurk trials (with correct voicing category and C=1 causal responses) according to their perceptual confidence levels separately for ‘B/P’ (Figure 7 A, D), ‘D/T (Figure 7 B, E) and ‘G/K’ (Figure 7 C, F) percepts. We assessed statistically whether the response fractions for different perceptual confidence levels differed between congruent and McGurk stimuli. This statistical analysis could only be applied to ‘D/T’ and ‘G/K’ because there were not enough McGurk trials with a ‘B/P’ response. While we observed a trend (p = 0.1) for lower perceptual confidence on McGurk trials with D/T percepts (Table 5 A), for G/K trials the perceptual confidence was not different between McGurk and congruent trials (Table 5 B). Next, we assessed whether observers assigned lower causal confidence levels to McGurk than congruent stimuli, while accounting for the perceptual confidence levels. Again, we observed a non-significant trend towards lower causal confidence for McGurk relative to congruent trials only for ‘D/T’ (Table 5 C), but not for ‘G/K’ percepts (Table 5 D). Thus, observers integrate conflicting McGurk signals into auditory ‘G/K’ percepts that are associated with comparable perceptual and causal confidence as percepts on congruent ‘G/K’ trials. These results suggest that on those subsets of McGurk trials observers are no longer metacognitively aware of the conflicting ‘B/P’ phoneme and ‘G/K’ viseme stimuli.

**Figure 7.**
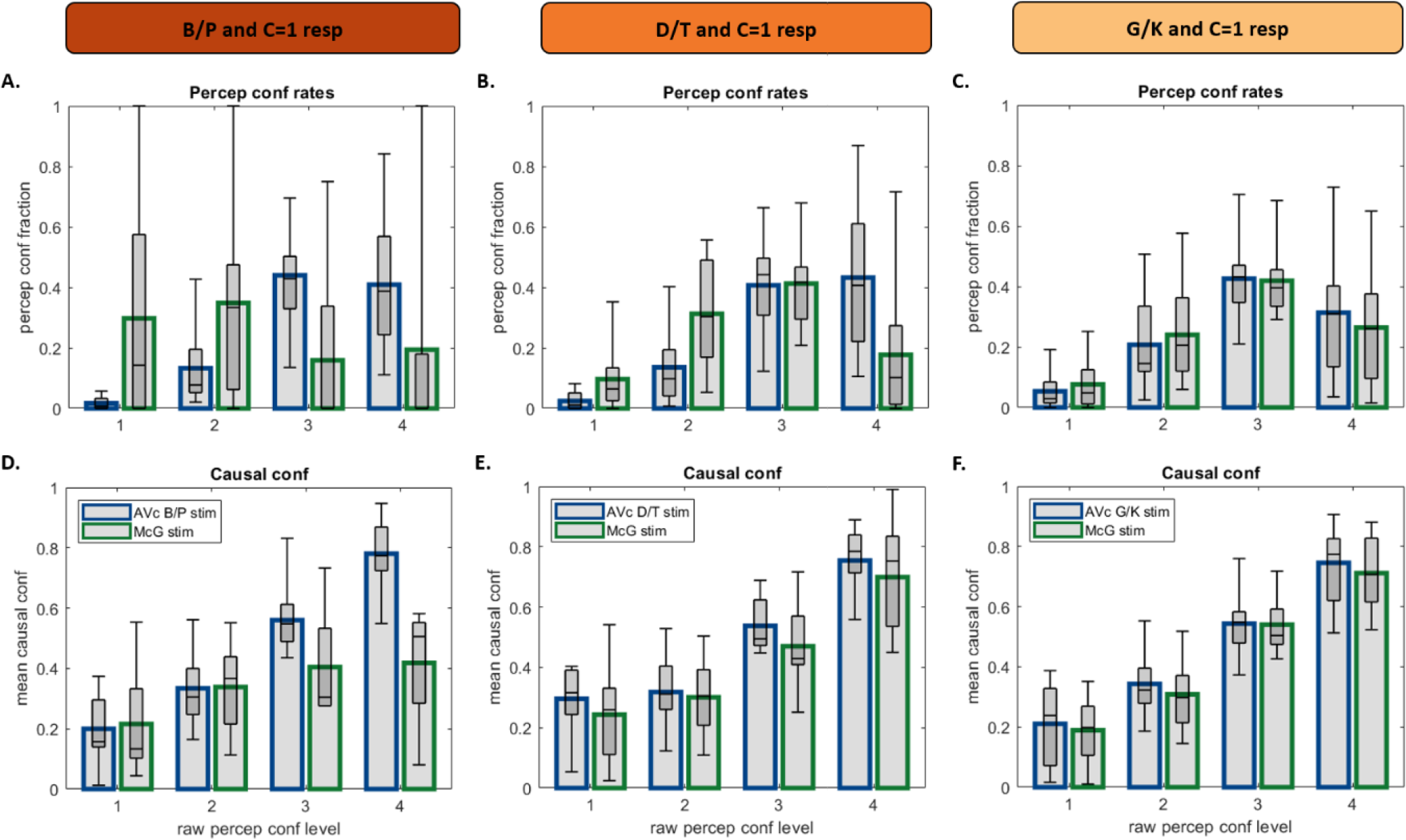
Perceptual and causal metamers. A, B, C. Response fractions over the four perceptual confidence levels for congruent stimuli with correct perceptual response and C=1 causal response (blue) and McGurk stimuli with correct perceptual voicing response and C=1 causal response (green) separately for ‘B/P’ (A), ‘D/T’ (B) and G/K (C) percepts. D, E, F. Causal confidence for the four perceptual confidence levels for congruent (blue) and McGurk (green) trials (as in A-C) separately for ‘B/P’ (A), ‘D/T’ (B) and G/K (C) percepts. In all figure components bar plots show across subjects’ means and box plots indicate across subjects’ median response fraction with first and third quartiles and with whiskers indicating the minimum and maximum subject values.

## Discussion

This study investigated how human observers form confidence judgments when presented with spoken syllables, articulatory lip movements or their congruent and McGurk combinations. In such multisensory information integration tasks, observers need to monitor two intimately related sorts of uncertainty: perceptual uncertainty about environmental properties (e.g. syllable’s first letter) and causal uncertainty about whether signals come from common or independent sources. Our results demonstrate that human observers form meaningful perceptual and causal confidence judgments that are qualitatively in line with the principles of Bayesian causal inference.

A wealth of research has shown that human observers integrate audiovisual signals from common sources weighted by their relative reliabilities into more precise percepts [38-41]. Sensory integration reduces observers’ uncertainty about the current state of the world. In our study, auditory and visual senses provide both redundant and complementary information about syllables [42]. The auditory sense facilitates the discrimination between voiced and unvoiced consonants (e.g. ‘B/G/D’ vs ‘P/K/T’) that is left ambiguous by the visual sense alone. Further, speech signals and the articulatory lip movements together inform about the place of articulation (e.g. ‘P/B’ vs. ‘D/T’ vs. ‘G/K’). Unsurprisingly, observers benefit substantially from audiovisual integration. They show superior syllable categorization accuracy and higher perceptual confidence on audiovisual congruent relative to unisensory trials (Figure 3). Only on less than 10% of the audiovisual congruent trials did observers miscategorize the syllables. As evidence for observers’ metacognitive sensitivity, they assigned lower levels of confidence to perceptual categorization errors relative to their correct responses [11]. Moreover, these perceptual categorization errors also revealed a close relationship between perceptual and causal inference. Observers’ causal accuracy and confidence were greater, when observers categorized the syllable correctly (Figure 5 A, C). Conversely, observers’ perceptual accuracy and confidence were greater for trials with high than low causal confidence (Figure 6 A, C). As we will discuss in greater detail below, this positive relationship between perceptual and causal accuracy resp. confidence is consistent with Bayesian causal inference models, in which perceptual and causal inference arise interactively and are susceptible to shared sensory noise [18, 21].

McGurk trials provide additional insights into the formation of confidence judgments by introducing a small intersensory conflict along the ‘place of articulation’ dimension, i.e. by combining an auditory B/P with a visual G/K signal. Unisensory auditory B/P stimuli were predominantly perceived as ‘B/P’, but in approximately 20% of the trials as ‘D/T’ or ‘G/K’. Unisensory visual G/K signals were mainly perceived as ‘G/K’ and in 20% of the trials as ‘D/T’, but nearly never as ‘B/P’. This inter-sensory difference in the distribution over perceptual categories explains that McGurk combinations are mainly integrated into ‘D/T’ and ‘G/K’ percepts that are possible perceptual explanations for both auditory ‘B/P’ and visual ‘G/K’ signals (Figure 4). Moreover, the perceptual outcome on a McGurk trial is characteristically influenced by observers’ causal inference on that trial. Consistent with Bayesian causal inference models, observers perceive the first letter of the auditory syllable as a ‘D/T’ or ‘G/K’ particularly, when they infer that auditory and visual signals come from common sources and hence integrate them (Figure 5, Figure 8). The proportion of visual biased ‘G/K’ percepts and the perceptual confidence increases even further for common source judgments with high relative to low causal confidence (Figure 6 B, D). Conversely, observers reported a veridical ‘B/P’ percept, i.e. a percept unbiassed by the conflicting visual G/K signal, when they inferred that auditory and visual signals come from independent sources. Again, this proportion of ‘B/P’ percepts increased for high relative to low causal confidence. Moreover, for both common and independent source responses, perceptual and causal confidence were positively related: a greater causal confidence was generally associated with a greater perceptual confidence. In short, McGurk trials replicated the tight relationship between perceptual and causal inference / confidence over trials that we already observed for congruent trials.

**Figure 8.**
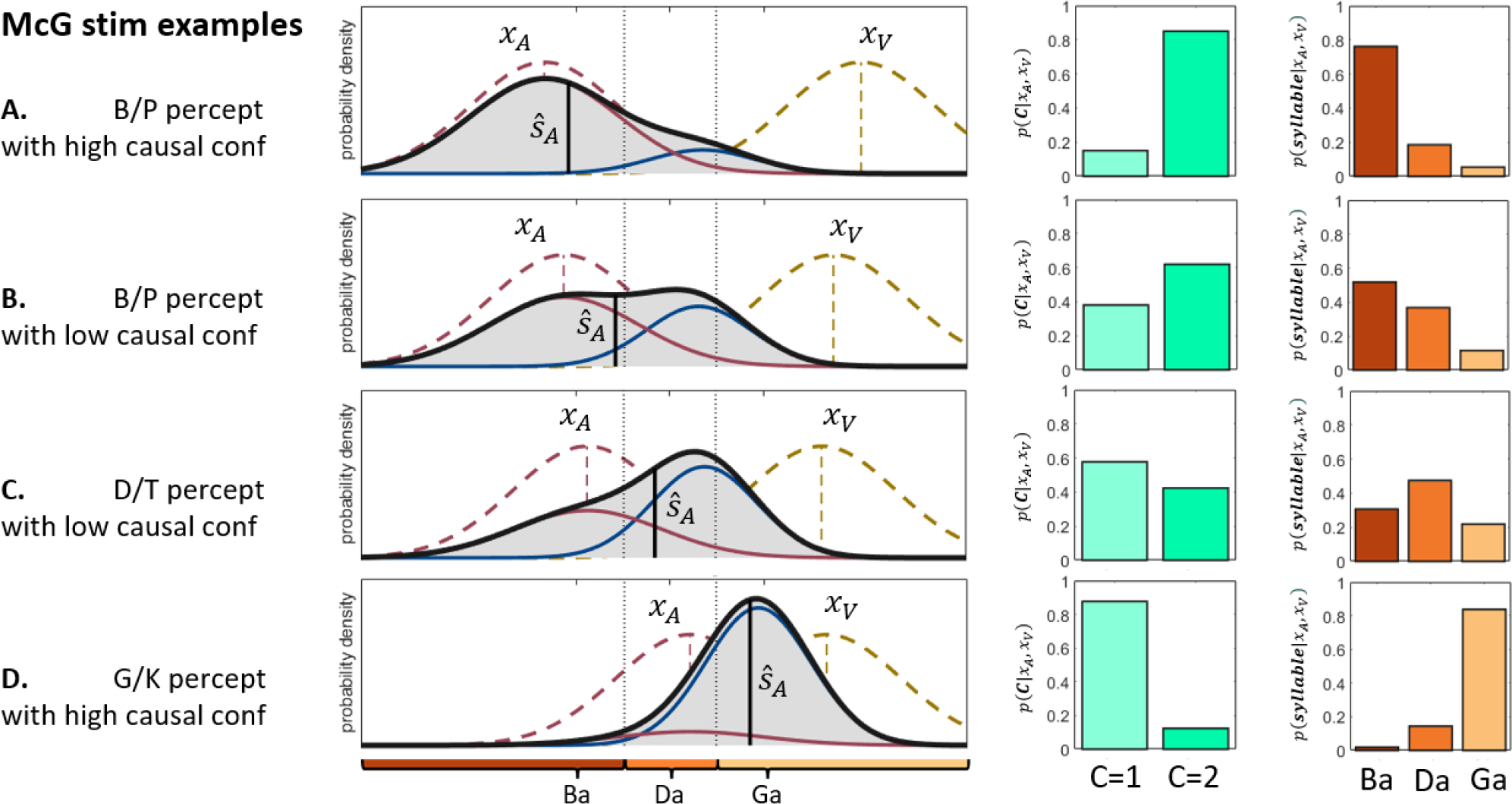
Simulation results of the Bayesian Causal Inference model. Each row shows the simulation results for a particular noisy auditory and visual signal pair that have been sampled from the generative model (see methods for details). The noisy auditory and visual signals were set to: A. *x*_*A*_ = -1.45; *x*_*V*_ = 2.20; B. *x*_*A*_ = -1.23; *x*_*V*_ = 1.88; C. *x*_*A*_ = -0.95; *x*_*V*_ = 1.76; D. *x*_*A*_ = 0.23; *x*_*V*_ = 1.80. For each sampled audiovisual signal pair, we show the likelihoods ℒ(*S*; *x*_*V*_) and ℒ(*S*; *x*_*A*_) (pink and brown dashed lines), the posterior distribution (solid black line) as a mixture of the full segregation *P*(*S*_*A*_|*x*_*A*_, *x*_*V*_, *C* = 2) (pink solid) and the fusion distributions *P*(*S*_*A*_|*x*_*A*_, *x*_*V*_, *C* = 1) (blue solid) weighted by the posterior probabilities over common and independent causes *P*(*C*|*x*_*A*_, *x*_*V*_) (green bar plots). The discrete posterior probabilities over ‘B/P’, ‘D/T’ and ‘G/K’ perceptual categories (orange bar plots) were obtained by integrating over the continuous posterior probability distribution between the respective category boundaries.

As shown in our simulations (Figure 8), this intimate relationship naturally arises in Bayesian causal inference models, because perceptual and causal inference are based on the same auditory and visual inputs that vary across trials because of sensory noise. Thus, when noisy auditory and visual signals are close together along the abstract ‘place of articulation dimension’ observers are likely to infer a common source and integrate audiovisual signals into a visual biased ‘G/K’ phoneme (Figure 8 bottom row). By contrast, an auditory dominant ‘B/P’ percept arises only, when the probability of independent sources is very high (Figure 8, top row).

While the Bayesian causal inference model can qualitatively explain the relationship between perceptual and causal decisions resp. confidence, our results cannot dissociate whether the brain forms Bayesian or approximate confidence estimates when exposed to multiple sensory signals under causal uncertainty. In these situations, the posterior probability distribution turns bimodal. Perceptual confidence may be related to a variety of quantities [43]. For instance, observers’ confidence judgments may reflect the posterior probability at the particular perceptual estimate or the entropy of the full bimodal posterior probability distribution [10, 44]. Alternatively, because observers perceive auditory speech signals categorically as ‘B/P’, ‘D/T’ or ‘G/K’, their perceptual confidence may be related to the posterior probabilities over the discrete response options rather than a continuous posterior distribution over a hypothesized place of articulation dimension. Further, in the discrete case, it is unclear whether observers’ perceptual confidence reflects Bayesian confidence, i.e. the posterior probability that a decision is correct or some other quantity. For example, in a three alternative visual categorization task observers’ perceptual confidence has recently been shown to reflect the difference in posterior probability between the two most likely options [45]. Because in our experimental paradigm observers made a choice amongst 6 options that were arranged in a two-dimensional ‘place of articulation’ x ‘manner of articulation’ space, it is likely that observers formed approximate or simple heuristic confidence judgments. Future research combining psychophysics with formal quantitative modelling is needed to dissociate between these different strategies to form confidence judgments.

McGurk trials provide critical insights into whether observers metacognitively monitor only the final integrated percept or whether they access the unisensory signals and underlying inference processes. An early intriguing study by Hillis et al. (2002) [32] has previously suggested that observers lose access to individual cues after integrating them within but not between the senses. In an oddity judgment task, conflicting visual- but not visuohaptic-cues were fused into perceptual metamers that were indistinguishable from the standard percepts derived from congruent cues. Following this rationale, we selected McGurk trials on which observers integrated the conflicting audiovisual signals into illusory ‘D/T’ and ‘G/K’ percepts and perceived a common cause. We then compared those to their corresponding perceptual and causal metamers of congruent trials, i.e. congruent audiovisual signals that elicited veridical ‘D/T’ and ‘G/K’ percepts and were perceived as coming from a common source (Figure 7) [30]. We reasoned that if observers move beyond the integrated percept and retain access to their unisensory ingredients, they should assign lower confidence to the conflicting McGurk signals than their congruent counterparts. Contrary to this conjecture we observed only non-significant trends towards lower perceptual and causal confidence for McGurk trials with ‘D/T’ percepts. For ‘G/K’ percepts, perceptual and causal confidence ratings were even more closely matched between congruent and McGurk stimuli (Figure 7). Thus, at least for visual biased ‘G/K’ percepts with common source judgments, observers obtained comparable confidence levels to perceptual and causal metamers that were unaffected by the underlying true causal structure. These results suggest that observers metacognitively monitor mainly the final integrated percept to form confidence judgments and do not access the unisensory signals or the true intersensory conflict when a common source has been inferred.

In conclusion, our results show that observers form meaningful causal and perceptual confidence estimates. Consistent with the principles of Bayesian causal inference, these two forms of uncertainty are closely related over trials with higher causal confidence typically associated with higher perceptual confidence. Further, when a common source of the sensory signals is inferred, confidence is directly informed by the final integrated percept with no or only very limited access to the unisensory signals and their true causal structure.

## Acknowledgments

We thank Sonal Patel for help with data acquisition. This research was funded by the ERC starting grant (‘multsens’).

## Supplementary Figures

**Figure S1:**
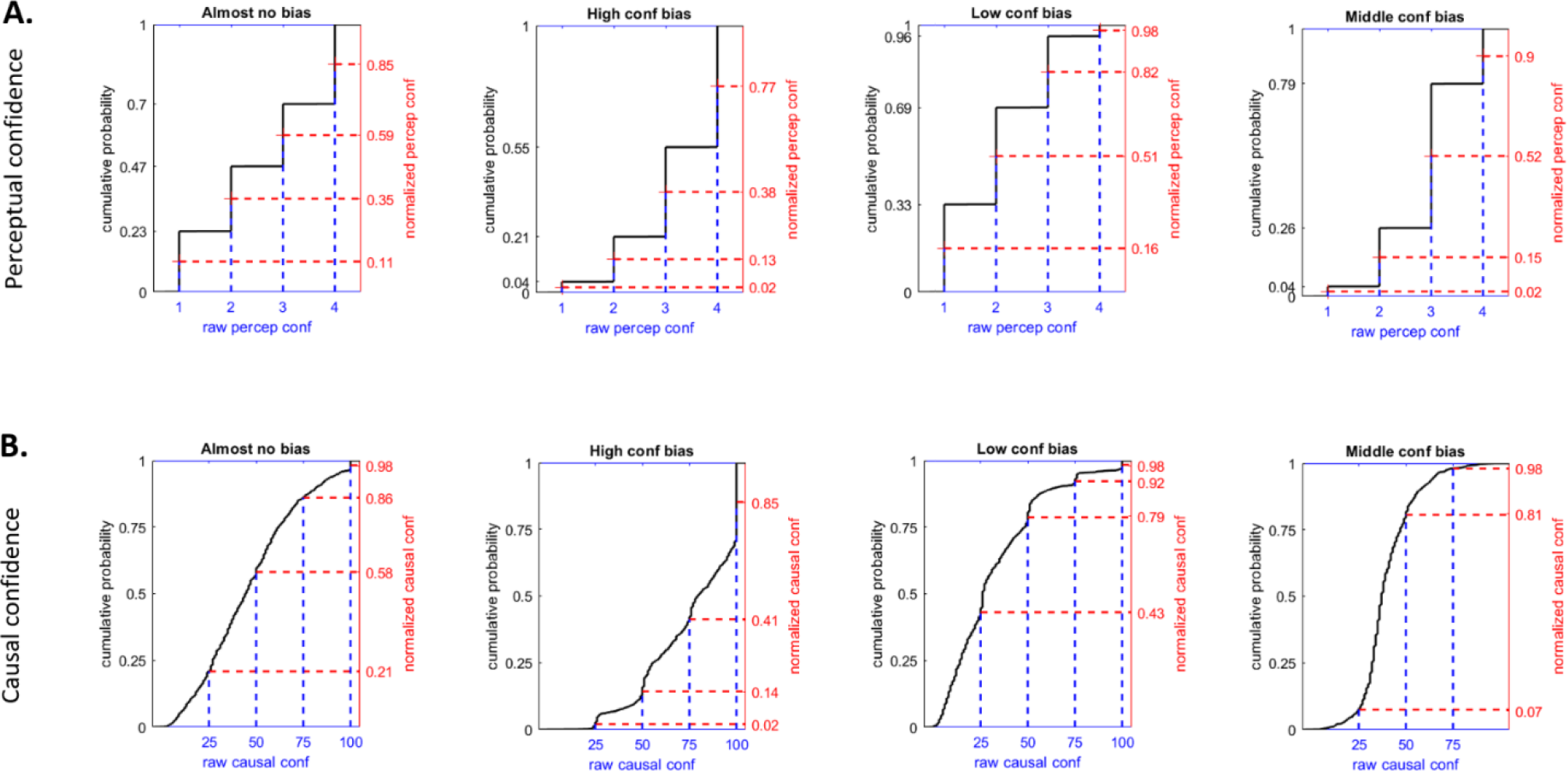
Confidence normalization. A. Cumulative distribution function (CDFs, across all trials, black) maps the raw perceptual confidence (blue, 4 levels) to the normalized perceptual confidence (red) for four exemplary subjects with variable confidence biases. B. Cumulative distribution function (CDFs, across all trials, black) maps the raw continuous causal confidence (blue) to their normalized causal confidence (red) for four exemplary subjects with variable confidence biases.

**Figure S2:**
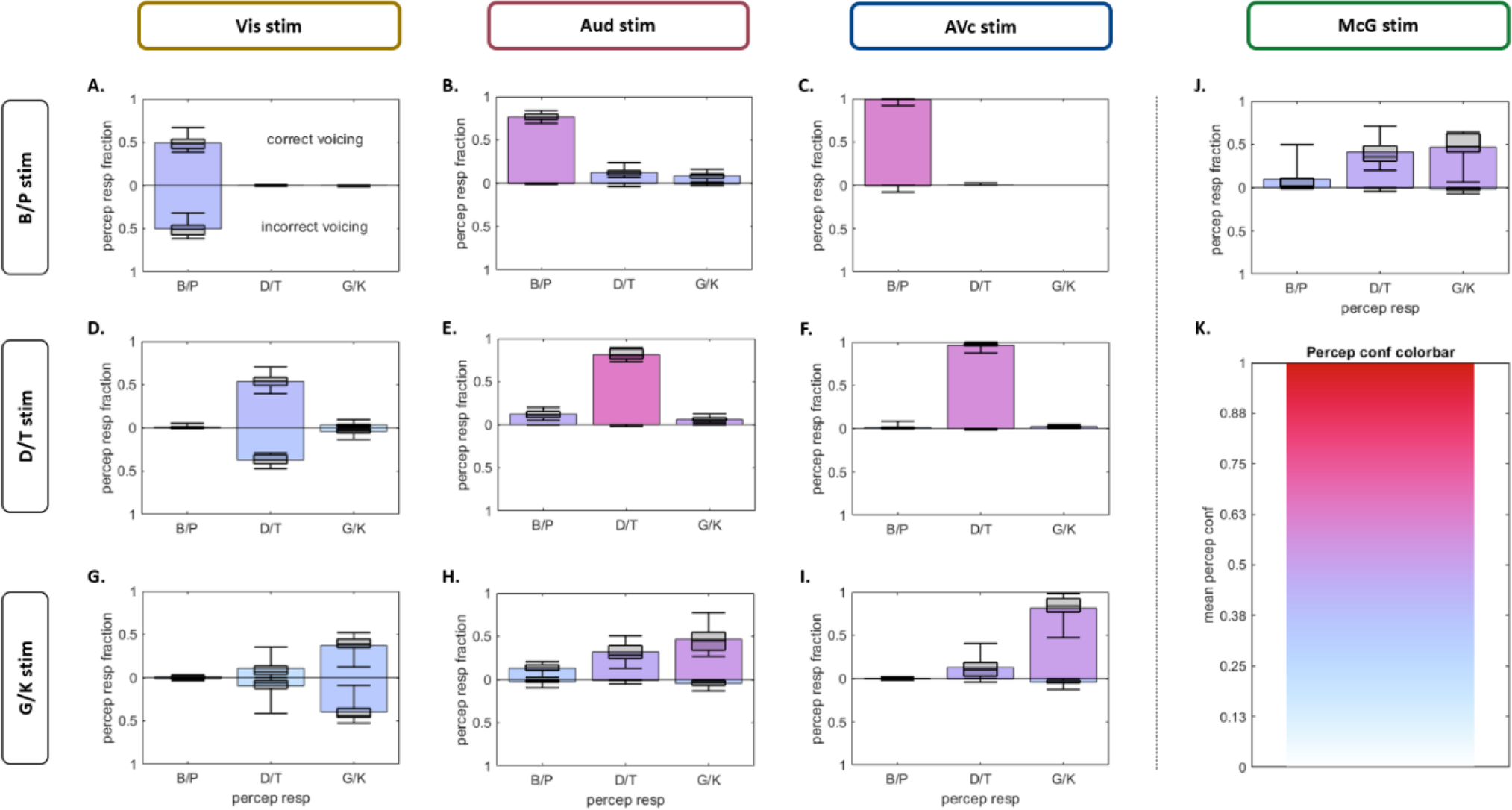
Perceptual response fractions and confidence. Response fractions over ‘B’, ‘P’, ‘D’, ‘T’, ‘G’, ‘K’ percepts across the three stimulus classes (rows): B/P stim, D/T stim and G/K stim and modalities (columns): Vis stim, Aud stim, AV congruent stim and McGurk stim. Each component figure shows the perceptual response fractions over place of articulation categories: ‘B/P’ vs. ‘D/T’ and ‘G/K’ along the x-axis. The voicing category (e.g. ‘B’ vs. ‘P’) is indicated along the y-axis. Incorrect voicing response fractions point downward in all subplots, whereas correct voicing response fractions point upward. Note that voicing errors were only common for unisensory Vis stimuli. The bar colors indicate the mean perceptual confidence for the respective response category, in accordance with the colorbar in panel K. Boxplots and whiskers: minimum subject, 25% quartile, median, 75% quartile, maximum subject.

## Supplementary Tables

**TABLE 1.** (see figure 3)

**1A. Accuracy**
| **1 x 3 rm-ANOVA** | **df** | **F** | **p** | **$\eta^2_p$** |
| --- | --- | --- | --- | --- |
| Stim type (Vis, Aud, AVc) | 2, 26 | 518 | < .001 | 0.98 |
| **Post hoc tests** |  | **t** | **p<sub>holm</sub>** | **Cohen's d** |
| Vis ≠ Aud |  | -12.9 | < .001 | -3.11 |
| Aud ≠ AVc |  | -19.1 | < .001 | -4.62 |

**1B. Percep conf for all responses**
| **1 x 3 rm-ANOVA** | **df** | **F** | **p** | **$\eta^2_p$** |
| --- | --- | --- | --- | --- |
| Stim type (Vis, Aud, AVc) | 1.40, 18.3 | 107 | < .001 | 0.89 |
| **Post hoc tests** |  | **t** | **p<sub>holm</sub>** | **Cohen's d** |
| Vis ≠ Aud |  | -9.47 | < .001 | -4.10 |
| Aud ≠ AVc |  | -4.95 | < .001 | -4.62 |

**1C. Percep conf for correct vs. incorrect**
| **2 x 3 rm-ANOVA** | **df** | **F** | **p** | **$\eta^2_p$** |
| --- | --- | --- | --- | --- |
| Percep resp type (correct, incorrect) | 1, 13 | 104 | < .001 | 0.89 |
| Stim type (Vis, Aud, AVc) | 1.43, 18.6 | 43.5 | < .001 | 0.77 |
| Percep resp type * Stim type | 2, 26 | 61.0 | < .001 | 0.82 |
| **Post hoc tests** |  | **t** | **p<sub>holm</sub>** | **Cohen's d** |
| Vis correct ≠ Vis incorrect |  | 2.20 | 0.145 | 0.75 |
| Aud correct ≠ Aud incorrect |  | 8.95 | < .001 | 3.06 |
| AVc correct ≠ AVc incorrect |  | 13.5 | < .001 | 4.60 |

**TABLE 2.** (see figure 4)

**2A. Accuracy for place of articulation**
| **Paired t-test** | **df** | **t** | **p** | **Cohen's d** |
| --- | --- | --- | --- | --- |
| Stim type (Aud B/P, Vis G/K) | 13 | -0.45 | 0.661 | -0.12 |

**2B. Entropy of percep resp distribution**
| **Paired t-test** | **df** | **t** | **p** | **Cohen's d** |
| --- | --- | --- | --- | --- |
| Stim type (Aud B/P, Vis G/K) | 13 | 2.53 | 0.025 | 0.68 |

**2C. Percep conf for B/P responses**
| **1 x 3 rm-ANOVA** | **df** | **F** | **p** | **$\eta^2_p$** |
| --- | --- | --- | --- | --- |
| Stim type (Aud B/P, Vis G/K, McG) | 2, 20 | 24.0 | < .001 | 0.71 |
| **Post hoc tests** |  | **t** | **$p_{holm}$** | **Cohen's d** |
| Aud B/P ≠ McG | | 2.78 | 0.012 | 1.19 |
| Vis G/K ≠ McG | | -4.11 | 0.001 | -1.76 |

**2D. Percep conf for D/T responses**
| **1 x 3 rm-ANOVA** | **df** | **F** | **p** | **$\eta^2_p$** |
| --- | --- | --- | --- | --- |
| Stim type (Aud B/P, Vis G/K, McG) | 2, 24 | 35.6 | < .001 | 0.75 |
| **Post hoc tests** |  | **t** | **$p_{holm}$** | **Cohen's d** |
| Aud B/P ≠ McG | | -2.74 | 0.011 | -1.02 |
| Vis G/K ≠ McG | | -8.28 | < .001 | -3.09 |

**2E. Percep conf for G/K responses**
| **1 x 3 rm-ANOVA** | **df** | **F** | **p** | **$\eta^2_p$** |
| --- | --- | --- | --- | --- |
| Stim type (Aud B/P, Vis G/K, McG) | 2, 26 | 27.1 | < .001 | 0.68 |
| **Post hoc tests** |  | **t** | **$p_{holm}$** | **Cohen's d** |
| Aud B/P ≠ McG | | -5.35 | < .001 | -2.04 |
| Vis G/K ≠ McG | | -7.06 | < .001 | -2.69 |

**TABLE 3.** (see figure 5)

**3A. C=1 resp rates for AVc stimuli**
| Paired t-test | df | t | p | Cohen's d |
| --- | --- | --- | --- | --- |
| Percep resp type (correct, incorrect) | 13 | 6.13 | < .001 | 1.64 |

**3B. C=1 resp rates for McG stimuli**
| 1 x 3 rm-ANOVA | df | F | p | $\eta^2_p$ |
| --- | --- | --- | --- | --- |
| Percep resp (B/P, D/T, G/K) | 1.35, 16.2 | 10.3 | 0.003 | 0.46 |
| Post hoc tests | | t | $p_{\text{holm}}$ | Cohen's d |
| B/P ≠ D/T | | -2.77 | 0.021 | -0.92 |
| D/T ≠ G/K | | -1.73 | 0.098 | -0.58 |

**3C. Causal conf for AVc stimuli**
| 2 x 2 rm-ANOVA | df | F | p | $\eta^2_p$ |
| --- | --- | --- | --- | --- |
| Percep resp type (correct, incorrect) | 1, 13 | 71.8 | < .001 | 0.85 |
| Causal resp (C=1, C=2) | 1, 13 | 5.02 | 0.043 | 0.28 |
| Percep resp type * Causal resp | 1, 13 | 53.6 | < .001 | 0.81 |
| Post hoc tests | | t | $p_{\text{holm}}$ | Cohen's d |
| correct C=1 ≠ correct C=2 | | 6.68 | < .001 | 2.34 |
| incorrect C=1 ≠ incorrect C=2 | | -3.45 | 0.004 | -1.21 |

**3D. Causal conf for McG stimuli**
| 3 x 2 rm-ANOVA | df | F | p | $\eta^2_p$ |
| --- | --- | --- | --- | --- |
| Percep resp (B/P, D/T, G/K) | 1.17, 10.6 | 1.29 | 0.291 | 0.13 |
| Causal resp (C=1, C=2) | 1, 9 | 0.72 | 0.419 | 0.07 |
| Percep resp * Causal resp | 2, 18 | 24.8 | < .001 | 0.73 |
| Post hoc tests | | t | $p_{\text{holm}}$ | Cohen's d |
| B/P C=1 ≠ B/P C=2 | | -4.32 | 0.003 | -1.56 |
| D/T C=1 ≠ D/T C=2 | | 2.37 | 0.234 | 0.85 |
| G/K C=1 ≠ G/K C=2 | | 3.74 | 0.012 | 1.35 |

**TABLE 4.** (see figure 6)

**4A. Accuracy for AVc stimuli**
| **2 x 2 rm-ANOVA** | **df** | **F** | **p** | **$\eta^2_p$** |
| --- | --- | --- | --- | --- |
| Causal resp (C=1, C=2) | 1, 13 | 60.0 | < .001 | 0.82 |
| Causal conf (low, high) | 1, 13 | 2.52 | 0.136 | 0.16 |
| Causal resp * Causal conf | 1, 13 | 11.3 | 0.005 | 0.47 |
| **Post hoc tests** |  | **t** | **p<sub>holm</sub>** | **Cohen's d** |
| C=1 low ≠ C=1 high | | -3.26 | 0.007 | -0.87 |
| C=1 low ≠ C=2 low | | 5.06 | < .001 | 1.61 |
| C=2 low ≠ C=2 high | | 0.69 | 0.497 | 0.19 |

**4B. B/P resp rates for McG stimuli**
| **2 x 2 rm-ANOVA** | **df** | **F** | **p** | **$\eta^2_p$** |
| --- | --- | --- | --- | --- |
| Causal resp (C=1, C=2) | 1, 13 | 6.92 | 0.021 | 0.35 |
| Causal conf (low, high) | 1, 13 | 5.04 | 0.043 | 0.28 |
| Causal resp * Causal conf | 1, 13 | 11.0 | 0.006 | 0.46 |
| **Post hoc tests** |  | **t** | **p<sub>holm</sub>** | **Cohen's d** |
| C=1 low ≠ C=1 high | | 1.38 | 0.348 | 0.18 |
| C=1 low ≠ C=2 low | | -1.42 | 0.348 | -0.48 |
| C=2 low ≠ C=2 high | | -4.00 | 0.003 | -0.53 |

**4C. G/K resp rates for McG stimuli**
| **2 x 2 rm-ANOVA** | **df** | **F** | **p** | **$\eta^2_p$** |
| --- | --- | --- | --- | --- |
| Causal resp (C=1, C=2) | 1, 13 | 16.6 | 0.001 | 0.56 |
| Causal conf (low, high) | 1, 13 | 3.19 | 0.097 | 0.20 |
| Causal resp * Causal conf | 1, 13 | 5.70 | 0.033 | 0.31 |
| **Post hoc tests** |  | **t** | **p<sub>holm</sub>** | **Cohen's d** |
| C=1 low ≠ C=1 high | | -2.97 | 0.025 | -0.55 |
| C=1 low ≠ C=2 low | | 2.74 | 0.027 | 0.93 |
| C=2 low ≠ C=2 high | | 0.58 | 0.108 | 0.57 |

**4D. Percep conf for AVc stimuli**
| **2 x 2 x 2 rm-ANOVA** | **df** | **F** | **p** | **$\eta^2_p$** |
| --- | --- | --- | --- | --- |
| Causal resp (C=1, C=2) | 1, 10 | 16.5 | 0.002 | 0.62 |
| Causal conf (low, high) | 1, 10 | 56.8 | < .001 | 0.85 |
| Percep resp type (correct, incorrect) | 1, 10 | 20.7 | 0.001 | 0.67 |
| Causal resp * Causal conf | 1, 10 | 15.0 | 0.003 | 0.60 |
| Causal resp * Percep resp type | 1, 10 | 0.27 | 0.618 | 0.03 |
| Causal conf * Percep resp type | 1, 10 | 2.65 | 0.135 | 0.21 |
| Causal resp * Causal conf * Percep resp type | 1, 10 | 0.04 | 0.843 | 0.00 |

**4E. Percep conf for McG stimuli**
| **2 x 2 x 3 rm-ANOVA** | **df** | **F** | **p** | **$\eta^2_p$** |
| --- | --- | --- | --- | --- |
| Causal resp (C=1, C=2) | 1, 5 | 0.03 | 0.865 | 0.01 |
| Causal conf (low, high) | 1, 5 | 13.3 | 0.015 | 0.73 |
| Percep resp (B/P, D/T, G/K) | 2, 10 | 9.35 | 0.005 | 0.65 |
| Causal resp * Causal conf | 1, 5 | 0.16 | 0.704 | 0.03 |
| Causal resp * Percep resp | 1.05, 5.24 | 2.84 | 0.150 | 0.36 |
| Causal conf * Percep resp | 2, 10 | 0.90 | 0.439 | 0.15 |
| Causal resp * Causal conf * Percep resp | 2, 10 | 2.94 | 0.099 | 0.37 |
| **Post hoc tests** |  | **t** | **p<sub>holm</sub>** | **Cohen's d** |
| B/P ≠ D/T | | -3.65 | 0.010 | -1.59 |
| B/P ≠ G/K | | -3.84 | 0.010 | -1.67 |
| D/T ≠ G/K | | -0.19 | 0.854 | -0.08 |

**TABLE 5.**
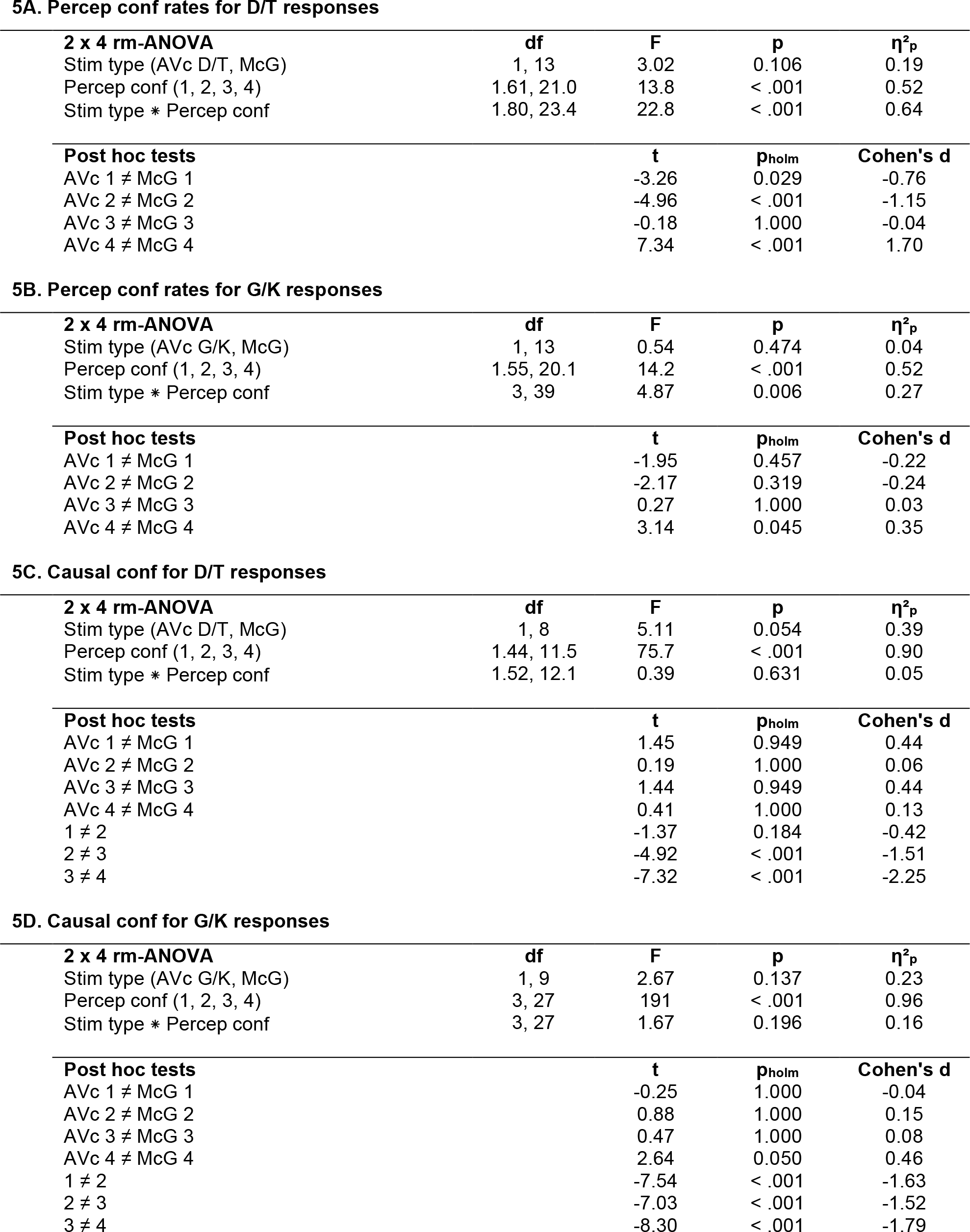
(see figure 7)

**5A. Percep conf rates for D/T responses**
| **2 x 4 rm-ANOVA** | **df** | **F** | **p** | **$\eta^2_p$** |
| --- | --- | --- | --- | --- |
| Stim type (AVc D/T, McG) | 1, 13 | 3.02 | 0.106 | 0.19 |
| Percep conf (1, 2, 3, 4) | 1.61, 21.0 | 13.8 | < .001 | 0.52 |
| Stim type * Percep conf | 1.80, 23.4 | 22.8 | < .001 | 0.64 |
| **Post hoc tests** |  | **t** | **p<sub>holm</sub>** | **Cohen's d** |
| AVc 1 ≠ McG 1 |  | -3.26 | 0.029 | -0.76 |
| AVc 2 ≠ McG 2 |  | -4.96 | < .001 | -1.15 |
| AVc 3 ≠ McG 3 |  | -0.18 | 1.000 | -0.04 |
| AVc 4 ≠ McG 4 |  | 7.34 | < .001 | 1.70 |

**5B. Percep conf rates for G/K responses**
| **2 x 4 rm-ANOVA** | **df** | **F** | **p** | **$\eta^2_p$** |
| --- | --- | --- | --- | --- |
| Stim type (AVc G/K, McG) | 1, 13 | 0.54 | 0.474 | 0.04 |
| Percep conf (1, 2, 3, 4) | 1.55, 20.1 | 14.2 | < .001 | 0.52 |
| Stim type * Percep conf | 3, 39 | 4.87 | 0.006 | 0.27 |
| **Post hoc tests** |  | **t** | **p<sub>holm</sub>** | **Cohen's d** |
| AVc 1 ≠ McG 1 |  | -1.95 | 0.457 | -0.22 |
| AVc 2 ≠ McG 2 |  | -2.17 | 0.319 | -0.24 |
| AVc 3 ≠ McG 3 |  | 0.27 | 1.000 | 0.03 |
| AVc 4 ≠ McG 4 |  | 3.14 | 0.045 | 0.35 |

**5C. Causal conf for D/T responses**
| **2 x 4 rm-ANOVA** | **df** | **F** | **p** | **$\eta^2_p$** |
| --- | --- | --- | --- | --- |
| Stim type (AVc D/T, McG) | 1, 8 | 5.11 | 0.054 | 0.39 |
| Percep conf (1, 2, 3, 4) | 1.44, 11.5 | 75.7 | < .001 | 0.90 |
| Stim type * Percep conf | 1.52, 12.1 | 0.39 | 0.631 | 0.05 |
| **Post hoc tests** |  | **t** | **p<sub>holm</sub>** | **Cohen's d** |
| AVc 1 ≠ McG 1 |  | 1.45 | 0.949 | 0.44 |
| AVc 2 ≠ McG 2 |  | 0.19 | 1.000 | 0.06 |
| AVc 3 ≠ McG 3 |  | 1.44 | 0.949 | 0.44 |
| AVc 4 ≠ McG 4 |  | 0.41 | 1.000 | 0.13 |
| 1 ≠ 2 |  | -1.37 | 0.184 | -0.42 |
| 2 ≠ 3 |  | -4.92 | < .001 | -1.51 |
| 3 ≠ 4 |  | -7.32 | < .001 | -2.25 |

**5D. Causal conf for G/K responses**
| **2 x 4 rm-ANOVA** | **df** | **F** | **p** | **$\eta^2_p$** |
| --- | --- | --- | --- | --- |
| Stim type (AVc G/K, McG) | 1, 9 | 2.67 | 0.137 | 0.23 |
| Percep conf (1, 2, 3, 4) | 3, 27 | 191 | < .001 | 0.96 |
| Stim type * Percep conf | 3, 27 | 1.67 | 0.196 | 0.16 |
| **Post hoc tests** |  | **t** | **p<sub>holm</sub>** | **Cohen's d** |
| AVc 1 ≠ McG 1 |  | -0.25 | 1.000 | -0.04 |
| AVc 2 ≠ McG 2 |  | 0.88 | 1.000 | 0.15 |
| AVc 3 ≠ McG 3 |  | 0.47 | 1.000 | 0.08 |
| AVc 4 ≠ McG 4 |  | 2.64 | 0.050 | 0.46 |
| 1 ≠ 2 |  | -7.54 | < .001 | -1.63 |
| 2 ≠ 3 |  | -7.03 | < .001 | -1.52 |
| 3 ≠ 4 |  | -8.30 | < .001 | -1.79 |

